# Functional connectivity dynamics reflect disability and multi-domain clinical impairment in patients with relapsing-remitting multiple sclerosis

**DOI:** 10.1101/2022.05.10.491171

**Authors:** Amy Romanello, Stephan Krohn, Nina von Schwanenflug, Claudia Chien, Judith Bellmann-Strobl, Klemens Ruprecht, Friedemann Paul, Carsten Finke

## Abstract

**Background:** Functional neuroimaging studies have revealed complex and heterogeneous patterns of aberrant functional connectivity (FC) in multiple sclerosis (MS), yet it remains unclear how time-resolved FC relates to variance in clinical disease severity.

**Objectives:** To characterize brain activity in MS patients with time-resolved FC analysis and explore the relationship between disease severity, multi-domain impairments, and altered network dynamics.

**Methods:** Resting-state functional MRI data were acquired from 101 MS patients and 101 age- and sex-matched healthy controls (HC). Dynamic FC analysis identified five connectivity states that were compared between HC and patients with high vs. low disability.

**Results:** Patients with higher disease severity exhibited a more widespread spatiotemporal pattern of altered FC and spent more time in a high-connectivity, low-occurrence state compared to patients with lower disease severity and HC. Depressive symptom severity was positively related to functional dynamics on global and network scales in patients, while fatigue and motor impairment were inversely related to frontoparietal network connectivity with the basal ganglia.

**Conclusions:** Time-resolved FC analysis uncovered alterations in network dynamics and clinical correlations that remained undetected with a static account of brain activity. Such time-varying approaches are thus crucial for disentangling the relationship between brain dynamics, disease severity, and symptoms in MS.

## INTRODUCTION

Identifying aberrant signatures of brain activity in multiple sclerosis (MS) is a key research target for improving our understanding of the complex interplay between symptom severity and functional network behavior. However, the characterization of common and clinically relevant functional alterations has been challenging, evidenced by a growing body of literature reporting complex, and occasionally, contradictory patterns of functional brain changes in MS.

Recent studies highlight subcortical brain regions as key players in the functional changes seen in MS: altered functional connectivity (FC) of the thalamus [1–3] and basal ganglia (BG) [4, 5] has been consistently linked to cognitive function and fatigue. In contrast, other studies have implicated both increased [6, 7] and opposingly, decreased default mode network (DMN) connectivity [8] in the relationship between functional alterations and cognitive impairment in MS.

Potential explanations for such heterogeneity include the temporally dynamic evolution of the disease, the ensuing variance in disease severity, its diverse cognitive and behavioral phenotypes as well as methodological variability across functional MRI (fMRI) studies. Furthermore, much previous research has been limited to a static account of brain activity in which FC is computed on signals over the entire scan duration. However, recent advances in time-resolved approaches – where FC is computed on a finer temporal scale and clustered into recurrent connectivity states – have made substantial contributions to identifying functional brain signatures in various states of health [9] and disease [10–12].

Such time-resolved techniques have recently highlighted reduced network centrality [13] and FC dynamics [14] in cognitively impaired MS patients, as well as subtype-specific altered FC and associations with cognitive and motor impairment [15]. While dynamic whole-brain connectivity studies thus represent a promising avenue to unravel the relationship between functional alterations and clinical outcomes, it remains unclear how altered network dynamics relate to disease severity and the multi-domain impairments commonly observed in MS.

Here, we characterize the spatiotemporal patterns of FC alterations in a large sample of patients with relapsing-remitting MS (RRMS). To this end, we investigate both static and dynamic FC alterations compared to healthy participants and relate these functional dynamics to disease severity and clinical impairment across multiple domains including cognitive, behavioral, and affective scores as well as structural brain measures.

## METHODS

### Participants and data acquisition

Patients were recruited from the outpatient clinic of the Clinical and Experimental Multiple Sclerosis Research Center at Charité - Universitätsmedizin Berlin from 2014-2018. At the time of their study visit, all patients fulfilled the McDonald criteria [16] for an RRMS diagnosis (Table 1). Patients did not have any comorbid neurological or psychiatric conditions. Healthy control participants (HC) were recruited using the Charité intranet and were without neurological or psychiatric conditions.

**Table 1.**
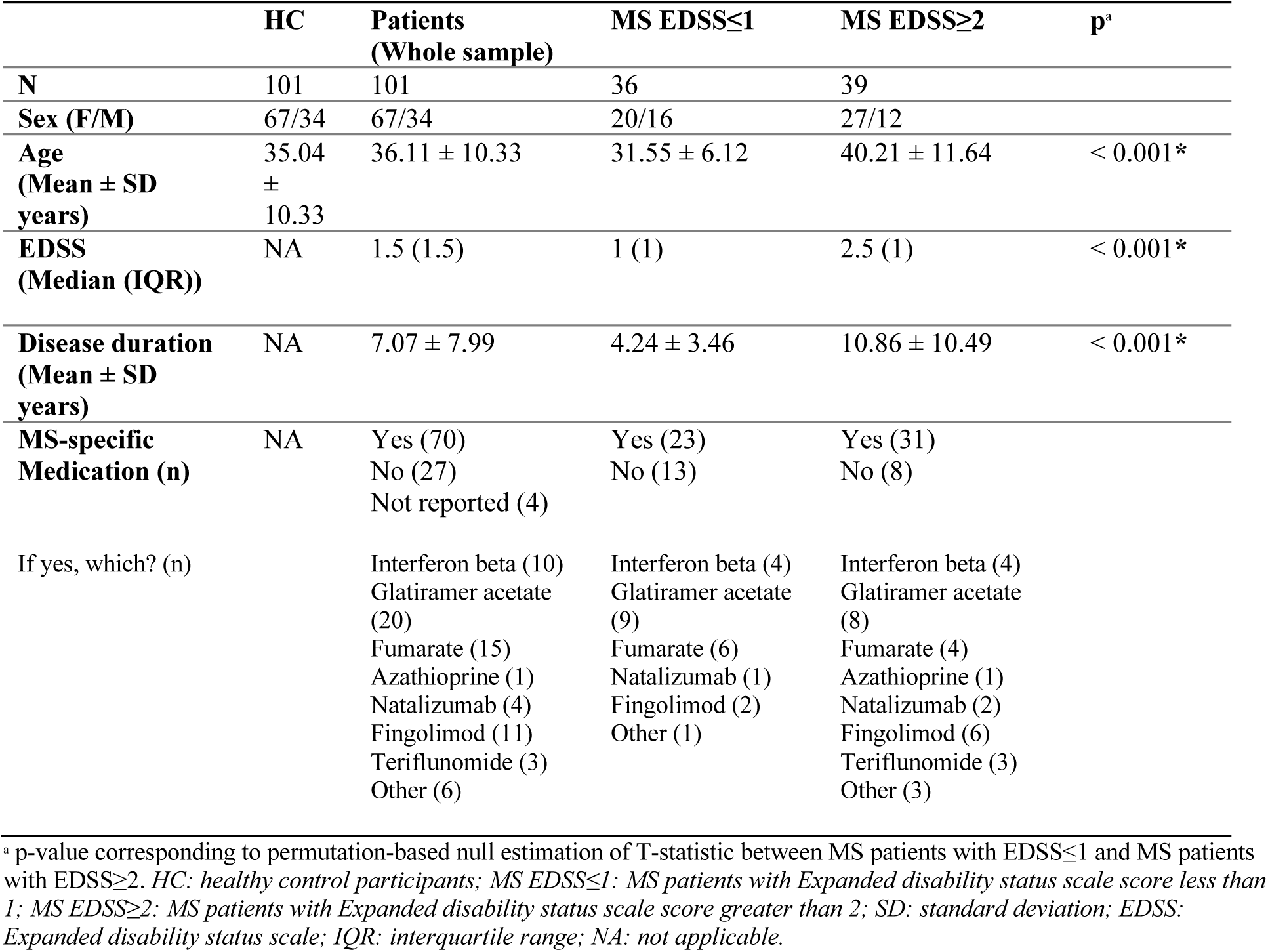
Demographic information for patients and healthy control participants.

Age- and sex-matching of patients and HCs was performed using a custom matching algorithm. Thus, 101 patients and 101 matched HCs were initially included; 92 patients had an Expanded Disability Status Scale (EDSS) rating from the date of scan acquisition. For the purposes of group comparisons, patients were split into subgroups of higher (EDSS≥2, n=39) and lower (EDSS≤1, n=36) disease severity using the upper and lower 30^th^ percentile threshold. Seventeen patients with an EDSS score of 1.5 were excluded. Patients with EDSS≥2 were significantly older and had a longer disease duration than patients with EDSS≤1 (Table 1). The final sample used for statistical analysis of group differences thus included 75 patients and 75 HCs. Additional details on the matching procedure and the patient cohort split are provided in supplemental material (SM) section 1.2.

MRI data were acquired at the Berlin Center for Advanced Neuroimaging at Charité-Universitätsmedizin Berlin, Germany, using a 20-channel head coil on a 3T Trim Trio scanner (Siemens, Erlangen, Germany). A 10-minute resting-state-fMRI (rs-fMRI) scan was collected using a repetition time (TR) of 2250ms (260 volumes, voxel size=3.4x3.4x3.4mm^3^). A T1-weighted structural scan was acquired using a magnetization-prepared rapid gradient echo (MPRAGE) sequence (1mm^3^ isotopic resolution). Additional details of MRI acquisition sequences are provided in SM section 1.1. All participants gave written informed consent, and the study was approved by the local ethics committee.

All patients also underwent an extensive battery of neuropsychological, motor, and visual acuity assessments. Test scores were grouped into 7 domains (cognition, motor, vision, depression, fatigue, as well as lesion load (total T2-Fluid-attenuated-inversion-recovery lesion volume), and total normalized brain volume. For each domain, a composite z-score was calculated (see SM section 1.3 for details and a list of measures assigned to each domain). To account for directionality differences, test scores for vision, cognition, and brain volume were re-coded by multiplying by - 1. Thus, composite z-scores represent impairment indices, where higher values signify worse performance or more severe symptoms.

### MRI data preprocessing

All analyses were performed with MATLAB (R2019b) and R (3.6.3). Preprocessing was performed using the CONN toolbox (version 18.b; [17]). Briefly, this included discarding five initial volumes of rs-fMRI scans, realignment, slice-timing correction, tissue segmentation, spatial normalization to Montreal Neurological Institute (MNI) space, and spatial smoothing.

T2-hyperintense brain lesion segmentation was performed manually using ITK-SNAP (by two MRI technicians with more than 10 years’ experience in MS research), and FSL cluster and fslstats were used to calculate lesion volumes and counts. Normalized brain volume was calculated from lesion-filled MPRAGE scans using FSL SIENAX.

### Group independent component analysis (GICA)

The spatial GICA feature of the Group ICA of fMRI toolbox (GIFT v4.0b, http://mialab.mrn.org/software/gift/index.html) was used to reduce whole-brain rs-fMRI data of all participants into 100 spatial group independent components. This method has been extensively described in [9, 18].

### Component rating and network assignment

Group ICs were manually classified as signal or noise based on criteria recommended in [19]. 47 group ICs were classified as “signal”, while 63 were discarded as “noise”. Signal components were assigned to the following cortical resting-state networks (RSN) according to [20]: ventral attention (vATT), dorsal attention (dATT), somatomotor (SMN), DMN, visual (VIS), and frontoparietal (FPN). Upon visual inspection, components overlapping subcortical (SC) and cerebellar (CB) brain regions were assigned to these groups for subsequent analyses. Additional details on preprocessing, GICA and component rating are provided in SM sections 1.4-6.

### Functional network analysis

Participants’ component time-courses underwent despiking, detrending, regression of realignment parameters and derivatives, and bandpass filtering (0.01-0.15Hz). Static FC was computed using pairwise Pearson’s correlations across all time-points. Dynamic FC was performed using sliding window correlations (as described in [9]), with a window size of 22 TR and a slide length of 1TR. Raw correlation coefficients were Fisher-Z transformed. K-means clustering was employed to extract distinct connectivity states using the “city-block” distance metric, as recommended in [21]. Based on convergence between the elbow criterion and Dunn’s Index [22], we identified five states. For reproducibility purposes, the numerical identity of the clusters was re-coded according to descending total occupancy. For each participant, the median over all dFC windows spent in a state was computed for further analyses. Additional details are provided in SM sections 1.7-8.

#### Dynamic metrics

We calculated the following additional measures of state dynamics: mean dwell time (the mean number of windows spent in a state upon entering it), fraction time (the total proportion of scan time spent in a state), transition frequency (the number of switches between each pair of states), and state stickiness (the number of times a participant remained in the same state over two consecutive windows).

State-wise average connectivity and modularity were calculated as measures of global connectivity and topology, respectively. Modularity was calculated using the community Louvain algorithm of the Brain Connectivity Toolbox (2019-03-03) [23]. Average connectivity was calculated as the global mean connectivity strength over all windows spent in that state. Additional details are provided in SM section 1.9.

### Network-wise overall connectivity

To explore the influence of differences in fraction time on global brain connectivity, we computed an additional “network-wise overall connectivity” (NWOC) metric. This was calculated by averaging across a participant’s total dFC data, resulting in one grand-average value across all connections and states, as well as 7 intra-NWOC values (not applicable to SC which had only one component), and 8 inter-NWOC values. Inter-NWOC values were computed as the overall connectivity between one RSN and the rest of the brain.

### Analysis of group differences

Between-group differences for static and dynamic FC were computed using a “non-parametric” two-sample T-test that estimates the null distribution by permuting group labels (see https://version.aalto.fi/gitlab/BML/bramila). For static FC and for each dFC state, differences were assessed between HC vs MS-EDSS≤1, HC vs MS-EDSS≥2, and MS-EDSS≤1 vs MS-EDSS≥2. Statistical significance was defined at an alpha level of 0.05 after false discovery rate (FDR) correction for multiple comparisons according to [24].

Group differences in dynamic metrics were assessed using a within-state approach. Linear regression was used to remove age-related variance. Using the model residuals, Kruskal-Wallis (KW) omnibus tests were performed. If the null hypothesis was rejected, post-hoc pairwise comparisons were performed using Dunn’s tests with FDR correction.

To test group differences in NWOC, an identical approach to the dynamic metric analyses was applied. In the case of group effects on inter-NWOC of (e.g., between the FPN and the rest of the brain), an additional analysis was performed between each pair of RSNs to identify which connections were driving the effect.

### Correlation analyses

Group differences in domain clinical outcome scores were evaluated using non-parametric permutation tests. Spearman’s partial correlations were calculated between patient’s FC values and clinical scores, controlling for age. FDR-correction was applied within state and domain. For correlations between dynamic metrics, NWOC, and clinical scores, Spearman’s correlations were computed using regression model residuals.

## RESULTS

### Clinical characterization

Patients with higher EDSS scores showed significantly worse outcomes across all clinical domains compared to patients with lower EDSS scores, including worse cognitive and motor performance, worse visual acuity, higher depression and fatigue symptoms, as well as higher lesion load and total brain atrophy (Table 2, Figure 1).

**Table 2.** Cognitive, behavioral, and structural brain composite Z-scores grouped by domain in patients. Table includes domain name, number of patients per group with complete data, group mean and standard deviation of computed Z-score, corrected p-value, and test statistic (T).

| Domain | N<br>(MS<br>EDSS≤1) | N<br>(MS<br>EDSS≥2) | Mean ± SD<br>(MS EDSS≤1) | Mean ± SD<br>(MS<br>EDSS≥2) | p<sub>FDR</sub> | T |
| --- | --- | --- | --- | --- | --- | --- |
| Vision<sup>a</sup> | 29/36 | 35/39 | -0.38 ± 0.46 | 0.31 ± 0.86 | < 0.001* | -3.987 |
| Motor<sup>b</sup> | 36/36 | 38/39 | -0.39 ± 0.51 | 0.40 ± 0.94 | < 0.001* | -4.534 |
| Fatigue<br>(FSS) | 33/36 | 36/39 | -0.63 ± 0.68 | 0.58 ± 0.87 | < 0.001* | -6.347 |
| Depression<br>(BDI-II) | 29/36 | 33/39 | -0.37 ± 0.75 | 0.32 ± 1.06 | 0.004* | -2.918 |
| Cognition<sup>c</sup> | 34/36 | 33/39 | -0.19 ± 0.66 | 0.17 ± 0.72 | 0.023* | -2.097 |
| Total brain<br>atrophy<sup>d</sup> | 33/36 | 32/39 | -0.28 ± 0.96 | 0.28 ± 0.94 | 0.014* | -2.345 |
| Lesion load<sup>e</sup> | 35/36 | 39/39 | -0.36 ± 0.43 | 0.32 ± 1.22 | < 0.001* | -3.232 |

<sup>a</sup> visual acuity composite z-score using ETDRS and Sloan tests; <sup>b</sup> motor performance composite z-score using 9-Hole Peg Test and Timed 25-foot Walk Test; <sup>c</sup> cognitive performance composite score using Selective Reminding Tests, Spatial Recall Tests, Symbol Digit Modality Test, Paced Auditory Serial Addition Test, and Word List Generation test; <sup>d</sup> negative total normalized brain volume; <sup>e</sup> T2-FLAIR total lesion volume; All clinical outcome z-scores and composites are coded such that higher values represent worse performance, higher symptom severity, or higher atrophy (see supplemental materials section 1.3 for additional details). *MS EDSS≤1*: MS patients with expanded disability status scale score less than or equal to 1; *MS EDSS≥2*: MS patients with expanded disability status scale score greater than or equal to 2; *FDR*: false discovery rate; *FSS*: Fatigue severity scale; *BDI-II*: Beck's depression inventory – second edition.

**Figure 1.**
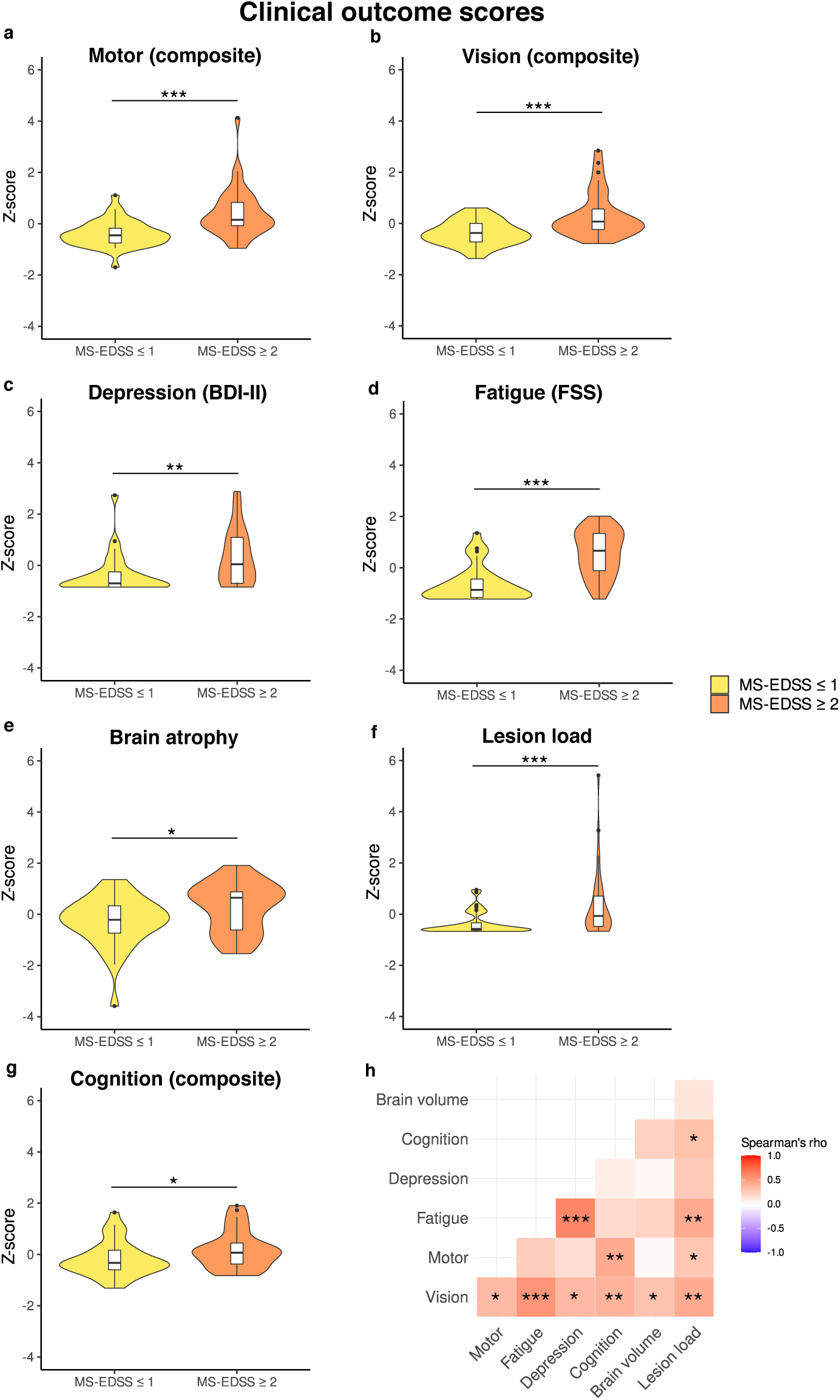
Between-group differences in clinical outcome z-scores across domains. Panels a-g: violin plots show distributions of clinical outcome scores in patients with lower (yellow) and higher (orange) severity; Motor, vision, and cognition domains are composite scores (see supplemental material section 1.3 for details). All clinical outcome z-scores and composites are coded such that higher values represent worse performance, higher symptom severity, or higher atrophy. Patients with higher disease severity in terms of Expanded disability status scale (EDSS) rating also showed higher impairment in all other domains. Panel h: correlation matrix where cells contain Spearman’s correlation coefficients between each pair of clinical outcome scores, computed across all patients. Asterisks indicate significance level: * = p_FDR_ < 0.05, ** = p_FDR_< 0.01, *** = p_FDR_ < 0.001.

### Functional network analysis

#### Static FC group differences

Group mean sFC and between-group comparisons are shown in Figure 2. Here, comparisons between patients with lower disease severity and matched HCs revealed decreased FC between the DMN and cerebellum in patients. Patients with higher disease severity showed a pronounced pattern of inter-network FC alterations compared to matched HCs, predominantly with FC increases between the DMN and FPN to other RSNs. In comparisons between both patient groups, more severely affected patients showed increased sFC between the FPN, SMN, and DMN and reduced connectivity between VIS-SMN and SC-dATT. However, these findings were not significant after FDR correction (Tables 3-5).

**Figure 2.**
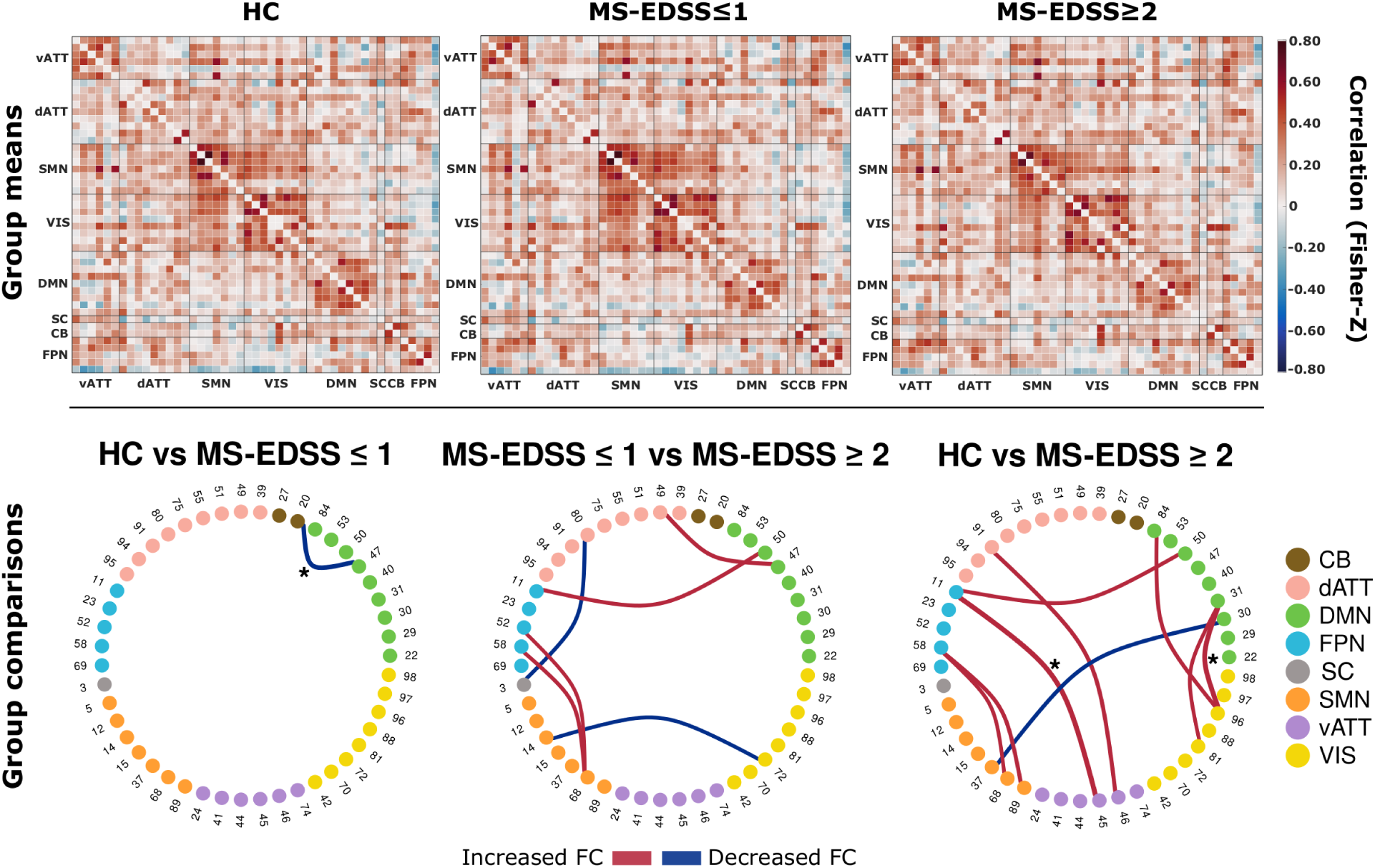
Whole-brain static functional connectivity matrices and between-group differences. Top: Mean static FC matrices computed by taking the average across all participants within a group. Pearson correlation coefficients were Fisher-Z transformed. Abbreviations on x- and y-axes indicate connections grouped by corresponding resting state networks. Matrix diagonals were zeroed for visualization purposes. Bottom: Between-group comparison results from the relevant permutation-based T-test. Group pairs used for comparison are denoted above each ring graph. Nodes and adjacent numbers identify the corresponding component (see supplemental material 1.6 for detailed component information). Nodes are grouped and color coded according to RSNs. Edge width maps the magnitude and direction of T-statistic. Edges were thresholded at an uncorrected p-value of 0.001. Asterisks indicate tests with a false discovery rate adjusted p-value < 0.05.

**Table 3.**
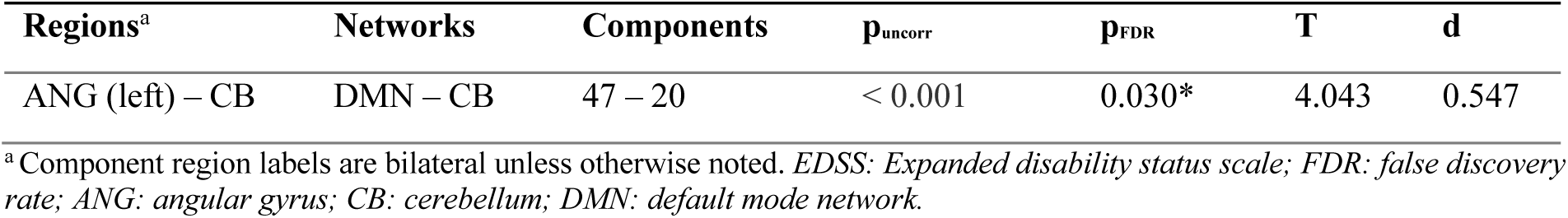
Static functional connectivity results for comparison between healthy participants and patients with EDSS≤1. Table includes brain region and network labels for the corresponding components, uncorrected p – values (p_uncorr._ ≤ 0.001), FDR – corrected p – values (p_FDR_), T – values and Cohen’s effect size (d).

**Table 4.** Static functional connectivity results for comparison between healthy participants and patients with EDSS≥2. Table includes brain region and network labels for corresponding components, uncorrected p-values (p_uncorr._), FDR-corrected p-values (p_FDR_), T-values and Cohen’s effect size (d).

| Regions<sup>a</sup> | Networks | Components | $p_{\text{uncorr.}}$ | $p_{\text{FDR}}$ | T | d |
| --- | --- | --- | --- | --- | --- | --- |
| Frontal pole – PCG | FPN – vATT | 11 – 45 | < 0.001 | < 0.001* | -4.429 | -0.831 |
| MFG – ACG | dATT – vATT | 91 – 46 | < 0.001 | 0.088 | -3.617 | -0.569 |
| STG – Precentral gyrus | DMN – SMN | 30 – 37 | < 0.001 | 0.088 | 3.513 | 0.646 |
| MFG (right) – SMA | FPN – SMN | 58 – 68 | < 0.001 | 0.088 | -3.286 | -0.622 |
| MFG (right) – Postcentral gyrus | FPN – SMN | 58 – 89 | < 0.001 | 0.065 | -3.564 | -0.621 |
| ANG (right) – Lingual gyrus | DMN – VIS | 31 – 81 | < 0.001 | 0.088 | -3.278 | -0.638 |
| ANG (right) – LOcc | DMN – VIS | 31 – 96 | < 0.001 | < 0.001* | -4.362 | -0.759 |
| Precuneus cortex – LOcc | DMN – VIS | 84 – 96 | < 0.001 | 0.088 | -3.481 | -0.713 |
| Frontal pole – Frontal pole | DMN – FPN | 50 – 11 | < 0.001 | 0.088 | -3.410 | -0.657 |

<sup>a</sup> Component region labels are bilateral unless otherwise noted. *EDSS*: Expanded disability status scale; *FDR*: false discovery rate; *PCG*: paracingulate gyrus; *MFG*: middle frontal gyrus; *ACG*: anterior cingulate gyrus; *STG*: superior temporal gyrus, posterior division; *SMA*: supplementary motor area; *ANG*: angular gyrus; *LOcc*: Lateral occipital cortex, superior division. *FPN*: fronto-parietal control network; *vATT*: ventral attention network; *dATT*: dorsal attention network; *DMN*: default mode network; *SMN*: somatomotor network; *VIS*: visual network.

**Table 5.** Static functional connectivity results for comparison between patients with EDSS≤1 and patients with EDSS≥2. Table includes brain region labels and network classification for corresponding components, uncorrected p-values (p_uncorr._), FDR-corrected p-values (p_FDR_), T-values and Cohen’s effect size (d).

| Regions<sup>a</sup> | Networks | Components | $p_{\text{uncorr.}}$ | $p_{\text{FDR}}$ | T | d |
| --- | --- | --- | --- | --- | --- | --- |
| ANG (left) – SMG | DMN – dATT | 47 – 49 | < 0.001 | 0.155 | -3.619 | -0.642 |
| BG – Superior parietal lobule (right) | SC – dATT | 3 – 80 | < 0.001 | 0.155 | 3.414 | 0.430 |
| Lingual gyrus – Precentral gyrus (left) | VIS – SMN | 72 – 14 | < 0.001 | 0.155 | 3.358 | 0.861 |
| MFG (left) – SMA | FPN – SMN | 52 – 68 | < 0.001 | 0.155 | -3.204 | -0.617 |
| MFG (right) – SMA | FPN – SMN | 58 – 68 | < 0.001 | 0.155 | -3.422 | -0.675 |
| Frontal pole – Frontal pole | FPN – DMN | 11 – 50 | < 0.001 | 0.155 | -3.476 | -0.724 |

<sup>a</sup> Component region labels are bilateral unless otherwise noted. *EDSS*: Expanded disability status scale; *FDR*: false discovery rate; *ANG*: angular gyrus; *SMG*: supramarginal gyrus; *BG*: basal ganglia; *MFG*: middle frontal gyrus; *SMA*: supplementary
*motor area. DMN: default mode network; dATT: dorsal attention network; SC: subcortical; VIS: visual network; SMN: somatomotor network; FPN: fronto-parietal control network.*

#### Dynamic FC states and group differences

Figure 3 shows the centroid position of each connectivity state and results of between-group comparisons. State 1 was the most frequently occurring and least globally connected state, while state 5 was the least frequently occurring state with highest global connectivity and lowest modularity. We observed an inverse relationship between the distributions of global average connectivity and modularity across states, as expected from previous network studies [25]. Additional state characteristics are shown in SM section 2.1.

**Figure 3.**
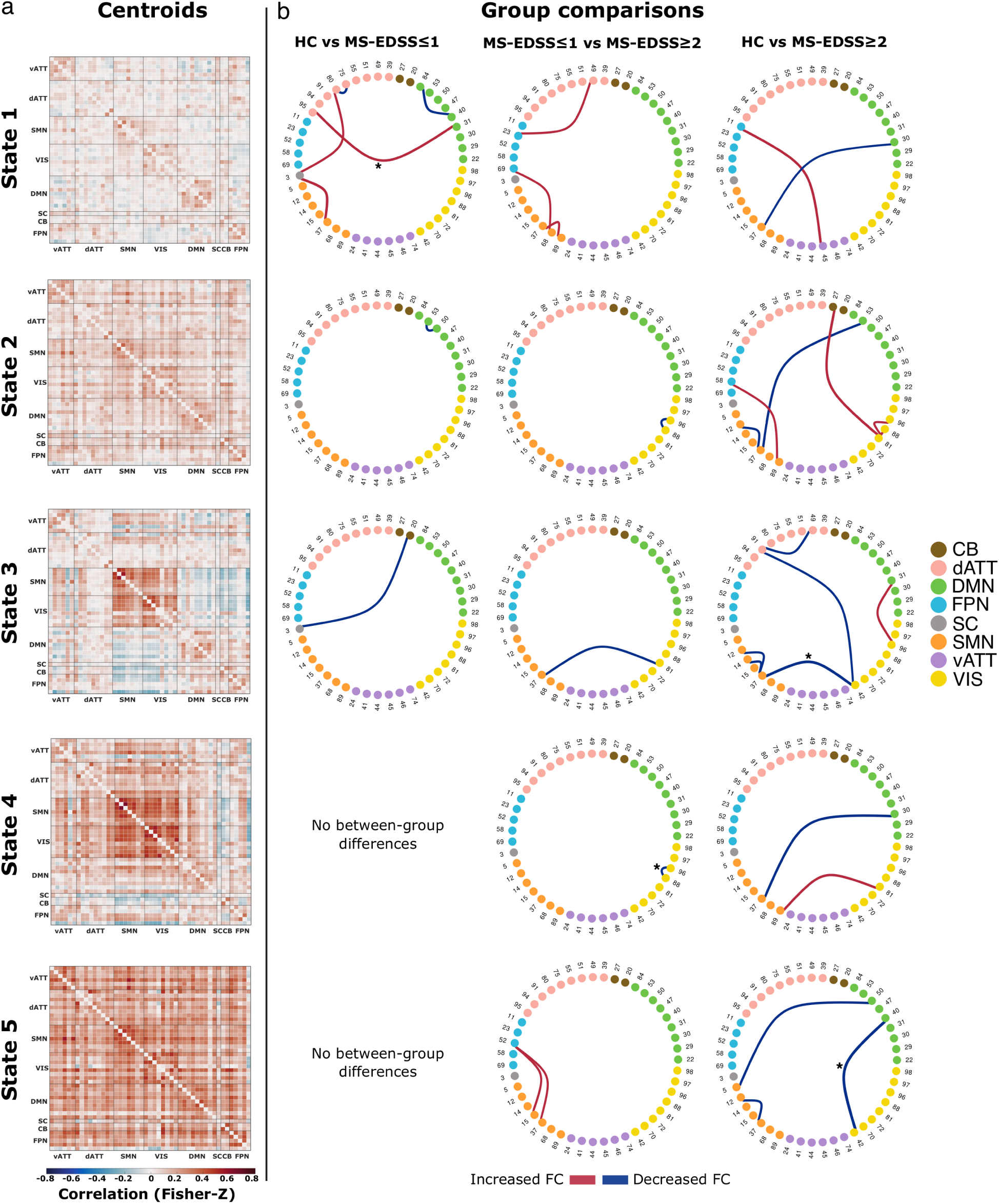
Whole-brain dynamic state centroids and between-group differences. Panel a: State centroid positions output from k-means clustering algorithm. Matrix diagonals were zeroed for visualization purposes. Panel b: Between-group comparison results from permutation-based T-tests. Group pairs used for comparisons are denoted above each column of ring graphs. Nodes and adjacent numbers identify the corresponding component (see supplemental material section 1.6 for detailed component information). Nodes are grouped and color coded according to their RSN assignment. Edge width maps the magnitude and direction of T-statistic. Edges were thresholded at an uncorrected p-value of 0.001. Asterisks indicate tests with an FDR-corrected p-value < 0.05.

State-wise between group comparisons revealed a widespread pattern of FC alterations. Compared to HCs, patients with EDSS≤1 had altered FC in three states – predominantly involving regions of the DMN and dATT. Interestingly, patients with higher severity showed a more spatially widespread and temporally persistent pattern of altered FC, implicating all connectivity states. Alterations were found in all five states and predominantly involved the VIS and SMN, as well as DMN, dATT, and FPN to a lesser extent. Comparisons between patient subgroups likewise revealed altered FC in all five states. In states 2-4, patients with higher disease severity exhibited reduced FC with the VIS. Additionally, patients with higher disease severity exhibited increased FC in states 1 and 5. This predominantly involved connections between the FPN-SMN and FPN-dATT. However, these increases were not significant after FDR correction (Tables 6-8).

**Table 6.** Dynamic functional connectivity results for comparison between healthy participants and patients with EDSS≤1. Table includes state labels, brain regions, and network labels for the corresponding components, uncorrected p-values (p_uncorr._), FDR-corrected p-values (p_FDR_), T-values and Cohen’s effect size (d).

| State | Regions <sup>a</sup> | Networks | Components | $p_{uncorr.}$ | $p_{FDR}$ | T | d |
| --- | --- | --- | --- | --- | --- | --- | --- |
| 1 | Superior parietal lobule (right) – Superior parietal lobule (left) | dATT – dATT | 80 – 75 | < 0.001 | 0.089 | 3.527 | 0.930 |
|  | BG – Superior parietal lobule (right) | SC – dATT | 3 – 80 | < 0.001 | 0.171 | -3.376 | -0.729 |
|  | ANG (right) – LOcc (left) | DMN – dATT | 31 – 95 | < 0.001 | 0.050* | -3.741 | -0.941 |
|  | BG – Precentral gyrus | SC – SMN | 3 – 37 | < 0.001 | 0.075 | -3.929 | -0.649 |
|  | Precuneus cortex – Precuneus cortex | DMN – DMN | 84 – 40 | < 0.001 | 0.089 | 3.593 | 0.645 |
| 2 | ANG (right) – Frontal pole | DMN – DMN | 53 – 50 | < 0.001 | 0.303 | 3.451 | 0.764 |
| 3 | CB – BG | CB – SC | 20 – 3 | < 0.001 | 0.501 | 3.456 | 0.709 |

<sup>a</sup> Component region labels are bilateral unless otherwise noted. *EDSS*: Expanded disability status scale; *FDR*: false discovery rate; *BG*: basal ganglia; *ANG*: angular gyrus; *LOcc*: Lateral occipital cortex, superior division; *ANG*: angular gyrus; *CB*: cerebellum; *dATT*: dorsal attention network; *SC*: subcortical; *DMN*: default mode network; *SMN*: somatomotor network.

**Table 7.**
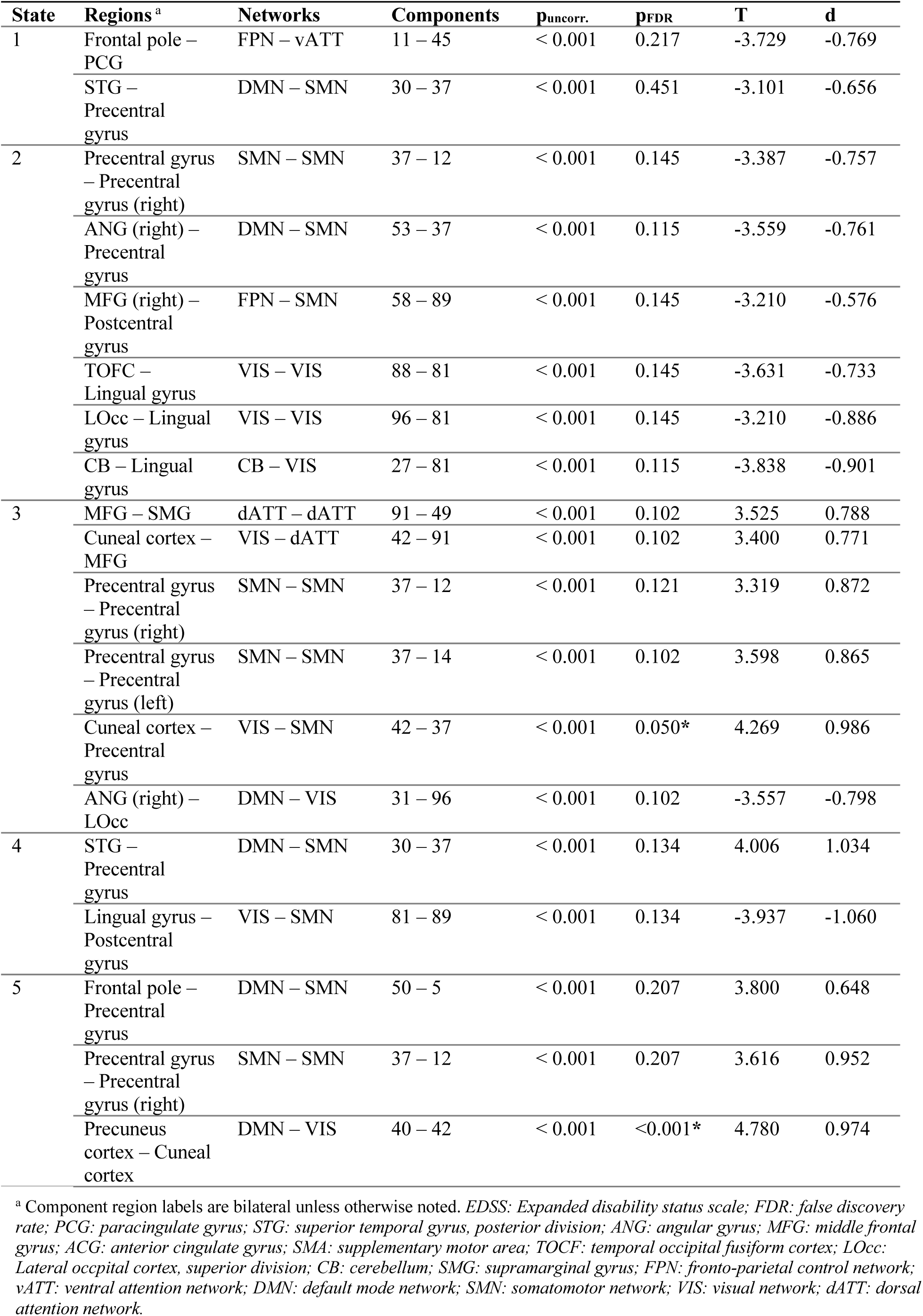
Dynamic functional connectivity results for comparison between healthy participants and patients with EDSS≥2. Table includes state labels, brain regions, and network labels for corresponding components, uncorrected p-values (p_uncorr_), FDR-corrected p-values (p_FDR_), T-values and Cohen’s effect size (d).

**Table 8.** Dynamic functional connectivity results for comparisons between patients with EDSS≤1 and patients with EDSS≥2. Table includes state labels, brain regions, and network labels for the corresponding components, uncorrected p-values (p_uncorr._), FDR-corrected p-values (p_FDR_), T-values and Cohen’s effect size (d).

| State | Regions <sup>a</sup> | Networks | Components | $p_{uncorr.}$ | $p_{FDR}$ | T | d |
| --- | --- | --- | --- | --- | --- | --- | --- |
| 1 | SMG – SMG | FPN – dATT | 23 – 49 | < 0.001 | 0.357 | -3.332 | -0.730 |
|  | Postcentral gyrus – Precentral gyrus | SMN – SMN | 89 – 37 | < 0.001 | 0.357 | -3.481 | -0.913 |
|  | LOcc – SMA | FPN – SMN | 69 – 68 | < 0.001 | 0.357 | -3.229 | -0.639 |
| 2 | LOcc – TOFC | VIS – VIS | 96 – 88 | < 0.001 | 0.270 | 3.626 | 0.747 |
| 3 | Lingual gyrus – Precentral gyrus | VIS – SMN | 81 – 37 | < 0.001 | 0.371 | 3.547 | 0.755 |
| 4 | LOcc – TOFC | VIS – VIS | 96 – 88 | < 0.001 | <0.001* | 4.485 | 0.911 |
| 5 | MFG (left) – Planum Temporale | FPN – SMN | 52 – 15 | < 0.001 | 0.368 | -3.500 | -0.787 |
|  | MFG (left) – Precentral gyrus | FPN – SMN | 52 – 37 | < 0.001 | 0.368 | -3.353 | -0.950 |

<sup>a</sup> Component region labels are bilateral unless otherwise noted. *EDSS*: Expanded disability status scale; *FDR*: false discovery rate; *SMG*: supramarginal gyrus; *LOcc*: Lateral occipital cortex, superior division; *SMA*: supplementary motor area; *TOCF*: temporal occipital fusiform cortex; *MFG*: middle frontal gyrus; *FPN*: fronto-parietal control network; *dATT*: dorsal attention network; *SMN*: somatomotor network; *VIS*: visual network.

### Dynamic metric group differences

No group effects were observed for average connectivity, modularity, or dwell time. In contrast, a significant group effect was present for state 5 stickiness as well as for transitions between states 3-5 and 4-5. Further analyses showed that this effect was due to differences in how often this state was visited. Accordingly, we observed a significant group effect on state 5 fraction time, such that more severely affected patients spent significantly more time in state 5 than both less severely affected patients and HCs (Table 9; Figure 4).

**Table 9.** Kruskal-Wallis rank sum test and post-hoc Dunn’s test for pairwise comparisons of group effects on state 5 fraction time residuals after age regression.

| Kruskal-Wallis test |  |  | Dunn's test |  |  |  |
| --- | --- | --- | --- | --- | --- | --- |
| $\chi^2$ | df | p | Comparisons | Z | $p_{uncorr.}$ | $p_{FDR}$ |
| 7.87 | 2 | 0.019* | HC – MS EDSS$\leq 1$ | 0.097 | 0.923 | 0.923 |
| | | | HC – MS EDSS$\geq 2$ | -2.612 | 0.009 | 0.027* |
| | | | MS EDSS$\leq 1$ – MS EDSS$\geq 2$ | -2.316 | 0.021 | 0.031* |

**Figure 4.**
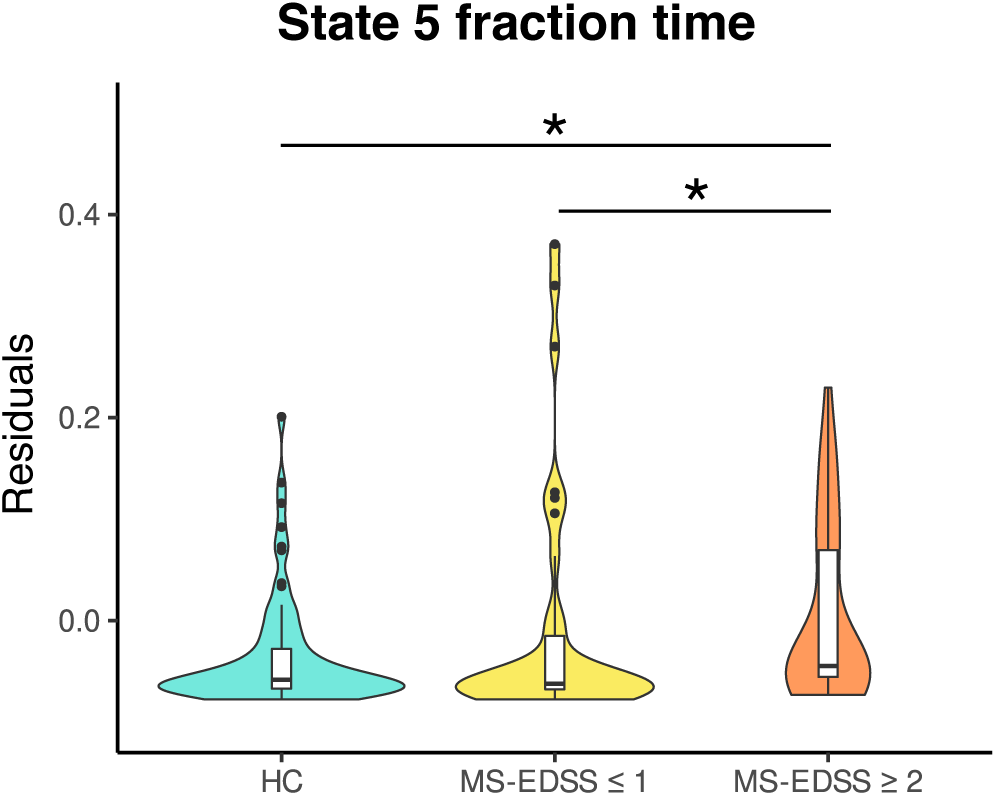
Group differences in state five fraction time. Violin plots show group-wise distributions of state five fraction time residuals after age-related variance was regressed out. Patients with higher disease severity (MS-EDSS≥2) spent significantly more time in state 5 compared to patients with lower disease severity (MS-EDSS≤1) and HCs. Outliers have been removed for visualization purposes. Asterisks indicate significance level: * = p_FDR_ < 0.05.

### Network-wise overall connectivity

Among all participants, state 5 had the highest average connectivity. Thus, if patients with EDSS≥2 spent more time in state 5, their average connectivity across all states may also be higher, as quantified by NWOC. On a global scale, we observed a gradual increase in NWOC from HC to more severely affected patients, but this effect was limited to a statistical trend. On the network scale, a significant group effect was found on inter-network connectivity between the FPN and all other networks, such that patients with EDSS≥2 had significantly higher FPN inter-NWOC than patients with EDSS≤1 and a trend in the same direction compared to HC (Table 10). We then further investigated which inter-network connections may drive this result.

**Table 10.**
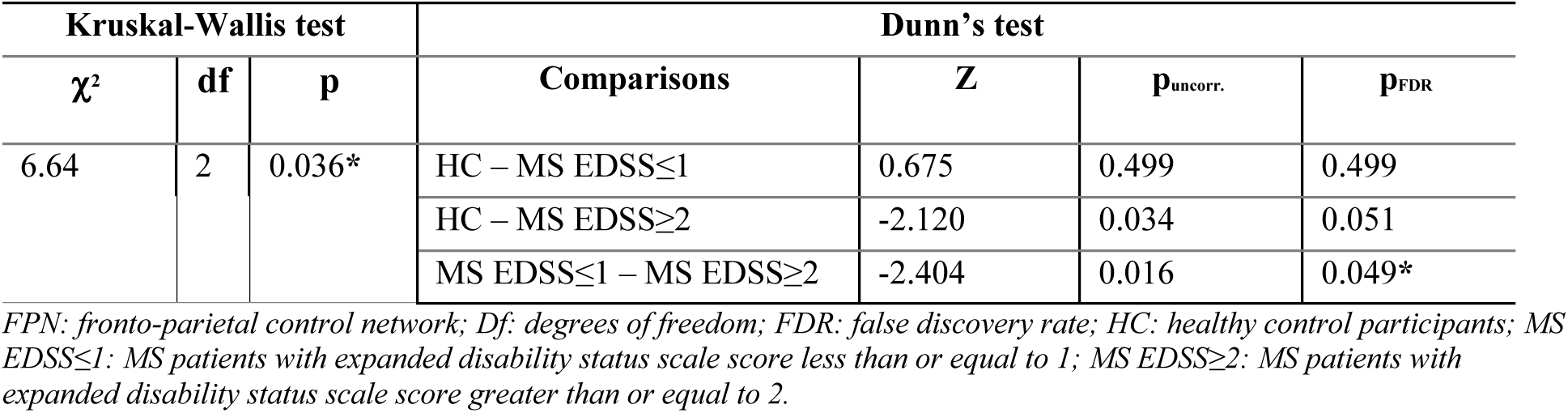
Kruskal-Wallis rank sum test and post-hoc Dunn’s test for pairwise comparisons of group effects on network wise overall connectivity between the FPN and rest of the brain.

Figure 5 shows that patients with higher disease severity had significantly higher FPN-SMN NWOC than both less severely affected patients and HCs, as well as significantly lower FPN-SC NWOC compared to less affected patients and a similar trend compared to HCs (Tables 11-12). Additionally, FPN-SC connectivity was negatively correlated with fatigue and motor impairment. Control analyses using static FC data to compute NWOC corroborated these findings. However, group effects on static NWOC between the FPN and the rest of the brain were not significant. Additional details are provided in SM section 2.2.

**Figure 5.**
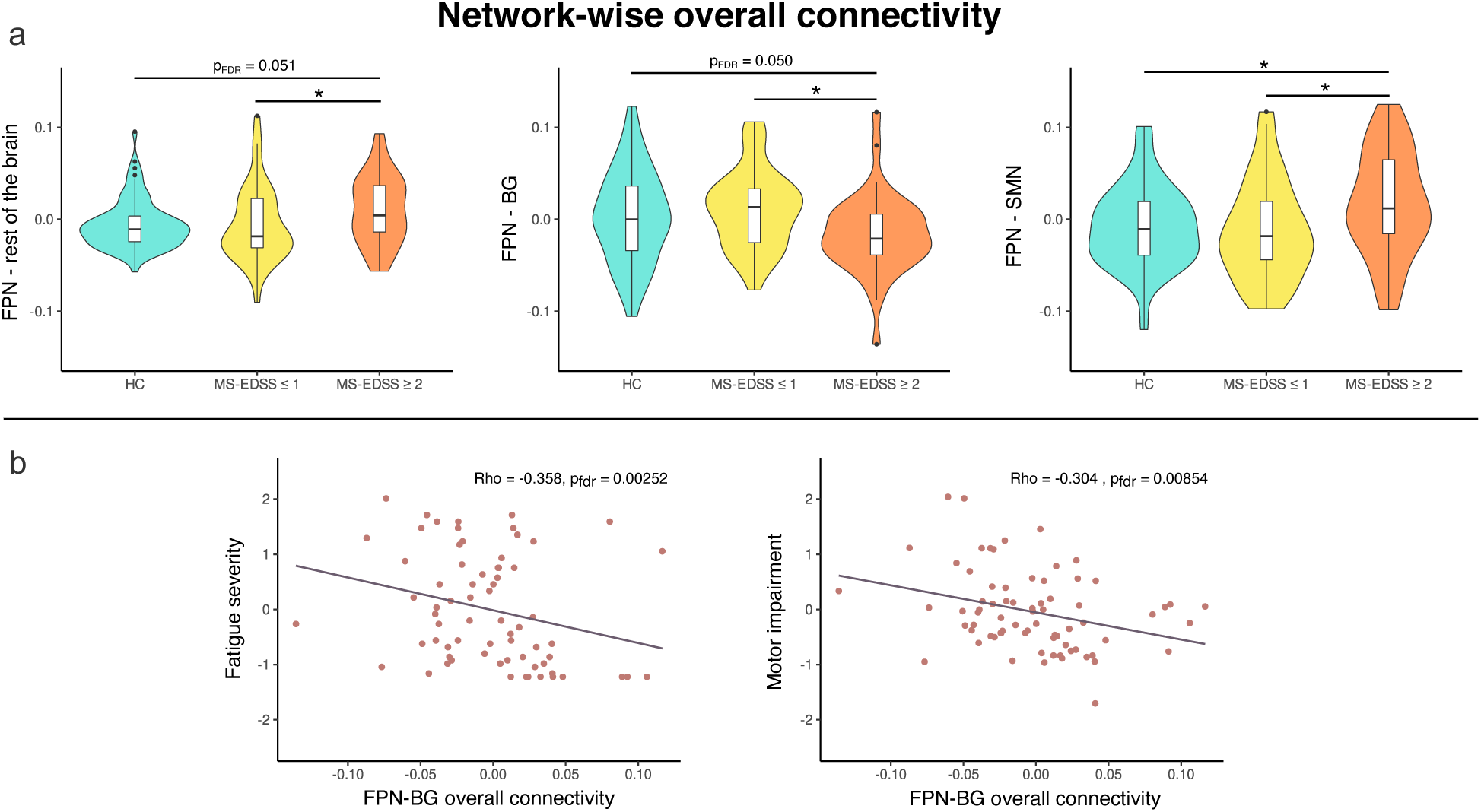
Frontoparietal network-wise overall connectivity group differences and correlations with clinical outcome scores. Panel a: (Left) Patients with higher disability severity have higher network-wise overall connectivity between the frontoparietal network (FPN) and the rest of the brain compared to patients with lower disability severity; (Middle) Patients with higher disability severity have lower network-wise overall connectivity between the frontoparietal network and basal ganglia (subcortical component) compared to patients with lower disability severity, with a similar trend compared to HCs; (Right) Patients with higher disability severity have higher network-wise overall connectivity between the frontoparietal network and somatomotor network (SMN) compared to patients with lower disability severity and HC. Panel b: Frontoparietal network-wise overall connectivity with the subcortex (basal ganglia) is negatively correlated with fatigue (left) and motor impairment (right); Asterisks denote significance level: * = p_FDR_ < 0.05.

**Table 11.** Kruskal-Wallis rank sum test and post-hoc Dunn’s test for pairwise comparisons of group effects on network wise overall connectivity between the FPN and SMN.

| Kruskal-Wallis test |  |  | Dunn's test |  |  |  |
| --- | --- | --- | --- | --- | --- | --- |
| $\chi^2$ | df | p | Comparisons | Z | $p_{uncorr.}$ | $p_{FDR}$ |
| 7.25 | 2 | 0.027* | HC – MS EDSS≤1 | 0.588 | 0.556 | 0.556 |
| | | | HC – MS EDSS≥2 | -2.283 | 0.022 | 0.034* |
| | | | MS EDSS≤1 – MS EDSS≥2 | -2.466 | 0.014 | 0.041* |

FPN: fronto-parietal control network; SMN: somatomotor network; Df: degrees of freedom; FDR: false discovery rate; HC: healthy control participants; MS EDSS ≤ 1: MS patients with expanded disability status scale score less than or equal to 1; MS EDSS ≥ 2: MS patients with expanded disability status scale score greater than or equal to 2.

**Table 12.** Kruskal-Wallis rank sum test and post-hoc Dunn’s test for pairwise comparisons of group effects network wise overall connectivity between the FPN and SC.

| Kruskal-Wallis test |  |  | Dunn's test |  |  |  |
| --- | --- | --- | --- | --- | --- | --- |
| $\chi^2$ | df | p | Comparisons | Z | $p_{uncorr.}$ | $p_{FDR}$ |
| 7.52 | 2 | 0.023* | HC – MS EDSS≤1 | -0.927 | 0.354 | 0.354 |
| | | | HC – MS EDSS≥2 | 2.125 | 0.034 | 0.050 |
| | | | MS EDSS≤1 – MS EDSS≥2 | 2.628 | 0.008 | 0.026* |

FPN: fronto-parietal control network; SC: subcortex; Df: degrees of freedom; FDR: false discovery rate; HC: healthy control participants; MS EDSS ≤ 1: MS patients with expanded disability status scale score less than or equal to 1; MS EDSS ≥ 2: MS patients with expanded disability status scale score greater than or equal to 2.

### Network-wise overall connectivity

#### Associations with clinical impairment

Figure 6 shows the results of correlation analyses between dFC, dynamic metrics, and clinical outcome scores that persisted when controlling for age. Lesion load was negatively correlated with both dATT-vATT and VIS-SMN connectivity in state 2. In contrast, lesion load was positively correlated with DMN-VIS connectivity in state 2. Depression was positively correlated with both DMN-vATT and dATT-vATT connectivity in state 3. Finally, total brain atrophy was positively correlated with DMN-VIS connectivity in state 3.

**Figure 6.**
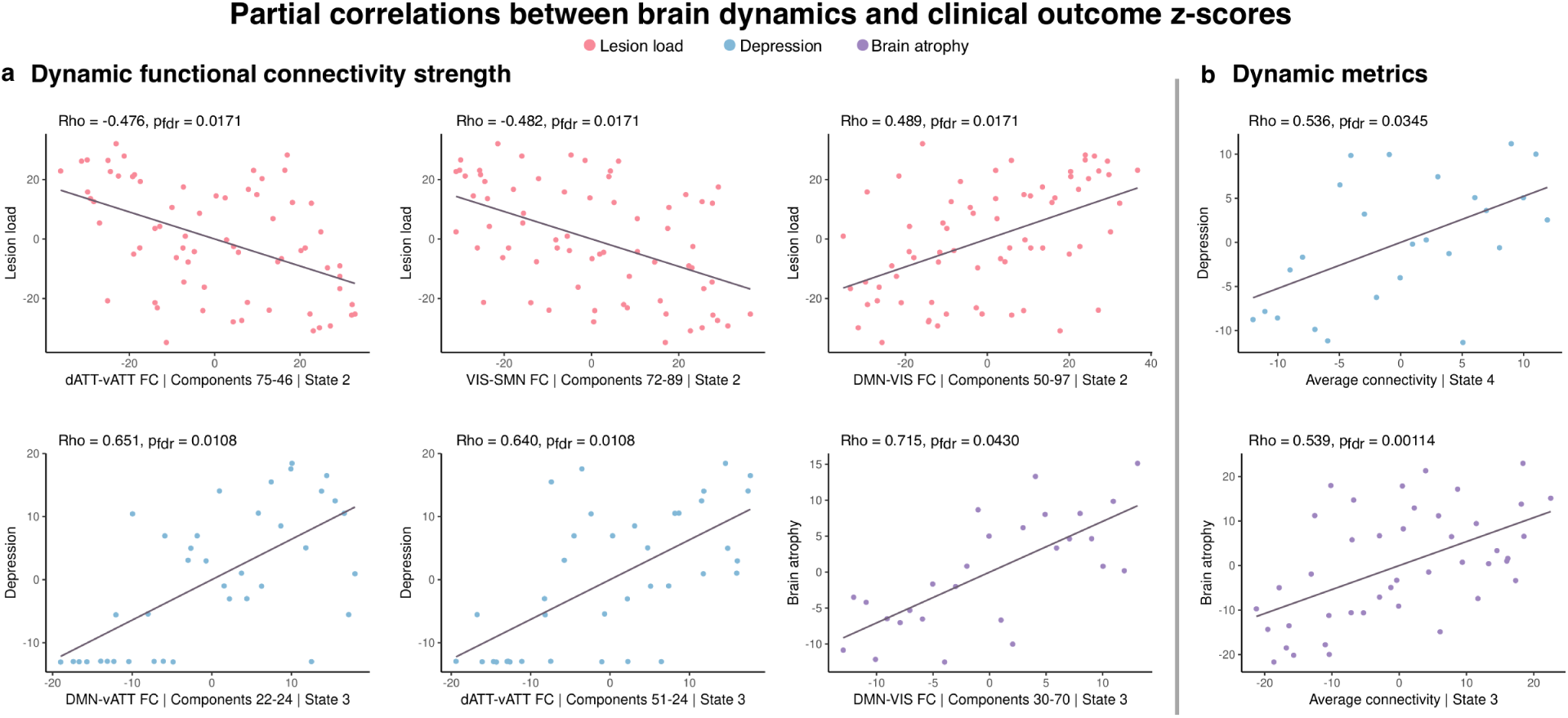
Partial correlations between dynamic functional connectivity, dynamic metrics, and clinical outcome z-scores. Panel a: Whole-brain dynamic functional connectivity values across patients were correlated with domain impairment scores. X-axes of subplots denote the resting-state network assignment, component numbers, and state in which the corresponding functional connection was significantly related to patient impairment indicated by the y-axis labels. Panel b: Dynamic metrics (x-axes) were correlated with domain impairment scores (y-axes). All correlations were performed using Spearman’s partial correlations controlling for age, and false discovery rate (FDR) correction was applied within state and domain. Values plotted correspond to variable residuals after rank-based age regression.

Correlation analyses between dynamic metrics and clinical scores revealed that depression severity was positively correlated with state 4 average connectivity, and brain atrophy was positively correlated with average connectivity of state 3.

Importantly, correlations between static FC strength and clinical scores were not significant. Similarly, no relationship was found between static global average connectivity and any domain clinical score. However, static modularity was positively correlated with brain atrophy (rho = 0.293, p_FDR_ = 0.037).

## DISCUSSION

In the present study, we show that functional brain dynamics in RRMS exhibit widespread alterations that vary with disease severity and reflect clinical outcomes across multiple symptom domains. With our whole-brain dynamic FC approach, we reveal distinct patterns of altered connectivity in patients with lower and higher disease severity as well as relationships with depressive symptoms and structural brain measures that remained undetected in static FC analyses. These findings highlight the advantages of time-resolved methods in studying functional network interactions and emphasize their essential role in understanding the complex relationships between brain activity, disease severity and clinical outcomes in MS.

We observed widespread bidirectional FC alterations across cortical and subcortical components in MS patients. While dynamic analyses revealed an intricate pattern of inter- and intra-network alterations, static FC analyses only detected inter-network changes. These observations suggest that sFC approaches may obscure transient intra-network connectivity changes that are observed only on the finer temporal scales of time-resolved analyses (i.e., seconds versus minutes).

Regarding the implicated brain systems, altered FC in the FPN and DMN has been reported in previous studies of patients with MS. For example, increased sFC of core FPN and DMN regions was associated with loss of cognitive efficiency [6]. Relatedly, another recent study reported increased connectivity between DMN and FPN to the rest of the brain in cognitively impaired patients with MS [7]. Static FC results of the present study support these findings, in that patients with higher disability showed increased FC between FPN, DMN and several other RSNs. However, age-controlled correlations between clinical scores and static FC did not yield any significant associations. With dynamic analyses, we observed consistent FPN and DMN involvement in FC alterations. While FPN connections to other RSNs were predominantly increased in patients with higher disease severity, DMN inter-network connectivity strength was mostly decreased in these patients.

We observed that increased depression symptoms were positively correlated with FC between the vATT-DMN and vATT-dATT in patients with MS. Importantly, the relevant DMN component corresponded to the bilateral hippocampus, which has been repeatedly implicated in depression [26] and shown to be affected in MS [27, 28]. In a study of hippocampal neuroinflammation, connectivity, and depression symptoms in MS patients, both positive and negative relationships between hippocampal resting-state connectivity and depression scores were reported [29]. While our results relating to hippocampal connectivity complement these findings, we furthermore identify a potential role of ventral and dorsal attention networks in MS-related depressive symptoms. This relationship between depression and FC increase is further supported by the positive association between depressive symptoms and global average connectivity in state 4. Taken together, these findings show that transient increases in FC, both globally and network-specific, are related to depressive symptom severity in patients with MS. The lack of this relationship in static connectivity analyses suggests that it may be specifically altered temporal brain dynamics that play a role in depressive symptoms in MS.

Fatigue and motor impairment are common features of MS. Several studies have reported inverse relationships between DMN-BG FC alterations and fatigue in MS. With a seed-based sFC approach, a negative relationship between fatigue severity and FC of BG with typical DMN structures was found [4, 30]. More recently, it was similarly reported with a dynamic approach that lower FC between the BG and DMN was associated with greater fatigue in patients with MS [5]. Here, we likewise observe the involvement of the BG, but instead implicate its connections to the FPN in MS-related fatigue. Additionally, we found a consistent clinical association of the FPN, with negative correlations between FPN-BG connectivity and motor impairment. These findings suggest a role of FPN-BG connectivity in clinical outcomes across multiple domains in MS.

Regarding the BG specifically, both white matter atrophy [31] and metabolic changes [32] have been observed in MS, and the role of BG in motor control is well-established [33]. However, FPN-BG interactions in patients with MS are less understood, as findings from previous studies commonly underscore DMN-BG connections. The FPN is typically regarded as a task-positive network involved in cognitive control and facilitating coordination between other brain networks [34]. Links between the FPN and various sub-regions of the BG in healthy participants are also well-documented [35]. With this context, the present findings suggest a maladaptive degradation of the FPN-BG coupling that is preserved in HCs and MS patients with lower disease severity but deteriorates in patients with higher disease severity. The observed inverse relationship between FPN-BG connectivity and motor and fatigue symptoms suggests that higher impairment in these domains is related to a decoupling of typical FPN and BG functional connections. Although these findings point to an exciting new brain target linking aberrant FC to clinical outcomes, further research is needed to characterize the precise role of BG connections with higher cognitive control networks and their associations with multi-domain impairment in MS.

### Limitations

Some limitations of the present work warrant mentioning. We analyzed a cross-sectional sample of data, which places an inherent limitation on investigating causal mechanisms of observed links between FC and clinical impairment. Additionally, neuropsychological and behavioral data were not sufficiently available for HCs and were thus not included in between-group analyses. This prevented us from making direct comparisons between patient subgroups and HCs when analyzing clinical outcome scores and limited the interpretation of impairment severity against normative values.

### Conclusions

These results show that connectivity alterations in MS are not homogeneous but depend on temporal network dynamics and disease severity, even in a sample with comparatively low overall disability status. Notably, the FPN was consistently implicated in both FC group differences and clinical associations to fatigue and motor impairment. Finally, dynamic analyses revealed both FC alterations and behavioral correlations that remained undetected in static analyses, highlighting that time-resolved accounts of brain activity disentangle the relationship between functional dynamics and clinical severity in MS.

