## Supplemental Materials for "Functional connectivity dynamics reflect disability and multi-domain clinical impairment in patients with relapsing-remitting multiple sclerosis"

1.0. Methods

1.1 MRI data acquisition parameters

Resting-state fMRI scans were collected using an echoplanar imaging sequence (TR = 2250ms, TE = 30ms, 260 volumes, matrix size = 64x64, 37 axial slices, slice thickness = 3.4mm, voxel size = $3.4x3.4x3.4mm^3$). High resolution T1-weighted structural scans were acquired using a magnetization-prepared rapid gradient echo (MPRAGE) sequence (matrix size = 240x240, 176 slices, $1mm^3$ isotropic resolution).

1.2. Participant selection and matching

Participants were selected from a large pool of RRMS patients and healthy control participants enrolled for an ongoing study and scanned using the same MRI sequence protocols. Framewise displacement (FD) according to [1] was calculated for each rs-fMRI scan and fifteen potential patient scans with mean FD > 0.55mm were excluded.

To optimize patient-control matching for age and sex differences, we implemented a matching algorithm using the MatchIt package (version 3.0.2, https://github.com/kosukeimai/MatchIt) for R (version 3.6.3)[1]. Propensity scores were estimated using a generalized linear model with the default logit link function. Matches were determined with greedy nearest neighbor matching, enforcing exact sex matching. To optimize the trade-off between pairwise age deviations and sample size, we applied a series of different caliper values (c=0.01 to 0.5; Figure 1). Calipers represent a liberality parameter defining the number of standard deviations of the distance measure within which to draw control units (https://r.iq.harvard.edu/docs/matchit/2.4-15/Additional_Arguments_f3.html). Accordingly, lower caliper values reflect stricter age optimization, leading to lower age deviations but also a smaller final study sample. We here chose the pivot point of the age-vs-sample plot as the optimal liberality parameter (arrow in Figure 1), as allowing for more pronounced age deviations in the matching after this point only yields small increases in the overall sample size while opting for stricter age matching results in a steep drop in sample size.

Finally, some of the control participants contributed more than one scan to the set of possible control matches (i.e., longitudinal data for these participants were available). To ensure that a particular control participant was only ever matched once, we randomly resampled the subset of longitudinal scans and repeated the matching procedure. Over 10000 repetitions, we thus found the best matches given the constraint that any control participant was uniquely matched to one patient.

This procedure yielded a final study population of n=202, with perfect sex matching (67 females, 34 males for both groups) and excellent age matching (healthy controls: mean age 35.04 ± 10.28 years [range 19.33 – 65.17]; patients: mean age 36.08 ± 10.27 years [range 19.32 – 62.52]; mean pairwise age deviation: 1.19 years; $t = -0.72$, $df = 200$, $p = 0.47$).

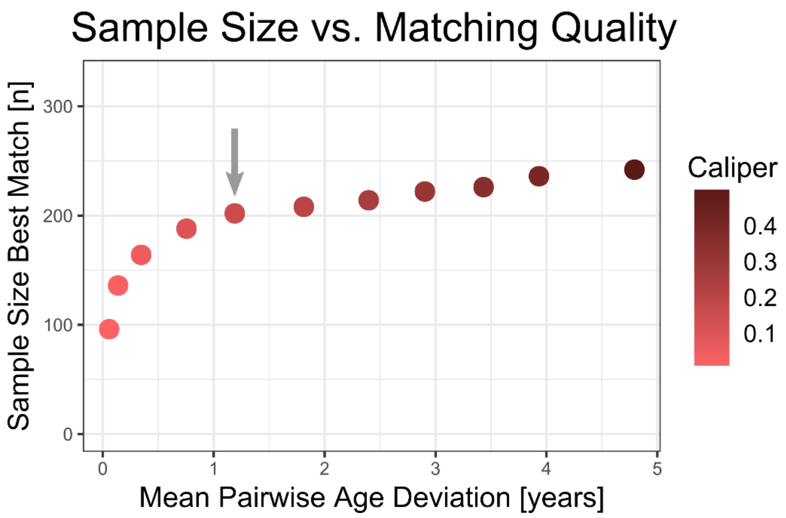


**Supplementary figure 1. Optimization of sample size and age deviation.** The plot shows the size of the study sample over the mean absolute age deviation of individual 1:1 matches as a function of the caliper values governing the strictness of age matching. The pivot point of this relationship is marked by an arrow and was chosen as the optimal liberality parameter for maximizing sample size while limiting age deviations.

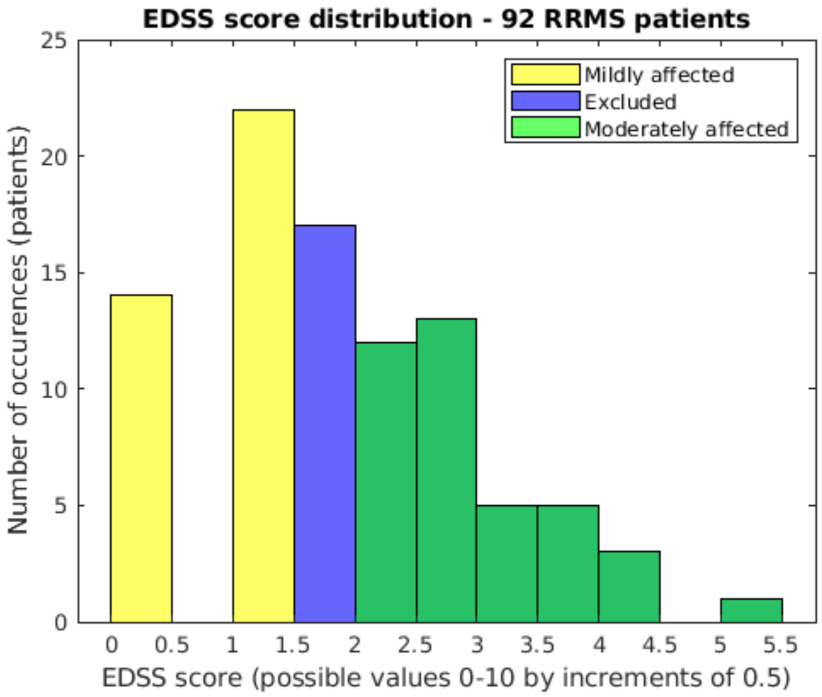


**Supplementary figure 2. Distribution of Expanded disability status scale (EDSS) scores for MS patients with data available from date of scan acquisition.** Following matching, 92 out of 101 patients had EDSS data available. Legend denotes patients grouped by disability severity, where "mildly affected" patients were those with an EDSS score in the lower 30% (MS-EDSS≤1) and "moderately affected" patients were those with an EDSS score in the upper 30% (MS-EDSS≥2) of the entire sample of (N = 92). 17 patients with EDSS score of 1.5 were excluded.

1.3. Domain impairment scores

Individuals test scores and structural brain measures were grouped into 7 domains (cognition, motor, vision, fatigue, depression, lesion load, and brain volume) for which Z-scores were subsequently calculated. The following procedure for computing domain composite z-scores was used: the summary or total score for each test was calculated across its items according to the respective test manual; the test's z-score was then computed across all patient scores; for domains which included multiple tests, a composite z-score was computed by averaging across all tests assigned to that domain. Thus, each patient ultimately had one value for each of the 7 domains.

**Supplementary table 1 –** Neuropsychological, cognitive, and behavioral questionnaires as well as structural brain measures assigned to each domain and included in impairment composite z-score calculations.

| Domain | Measures | Details |
|---|---|---|
| Visual acuity | ETDRS 100% high contrast | Right eye, decimal acuity |
| | | Left eye, decimal acuity |
| | | Right eye, number of letters correct |
| | | Left eye, number of letters correct |
| | ETDRS 10% medium contrast | Right eye, decimal acuity |
| | | Left eye, decimal acuity |
| | | Right eye, number of letters correct |
| | | Left eye, number of letters correct |
| | Sloan 2.5% low contrast | Right eye, decimal acuity |
| | | Left eye, decimal acuity |
| | | Right eye, number of letters correct |
| | | Left eye, number of letters correct |
| | Sloan 1.25% low contrast | Right eye, decimal acuity |
| | | Left eye, decimal acuity |
| | | Right eye, number of letters correct |
| | | Left eye, number of letters correct |
| Motor | Timed 25-Foot Walk Test (T25-FW) | |
| | Nine-Hole Peg Test (9-HPT) | |
| Fatigue | Fatigue severity scale (FSS) | |
| Depression | Beck's depression inventory II (BDI-II) | |
| Cognition | Selective reminding test – long-term storage (SRT-LR) | |
| | Selective reminding test – consistent long-term retrieval (SRT-CLTR) | |
| | Selective reminding test – delayed recall (SRT-DR) | |
| | Spatial recall test (SPAT) | |
| | Spatial recall test – delayed recall (SPAT-DR) | |
| | Symbol digit modality test (SDMT) | |
| | Paced auditory serial addition test (PASAT) | |
| | Word list generation test (WLG) | |
| Brain volume | Total normalized brain volume | |
| Lesion load | T2-FLAIR total lesion volume | |

ETDRS: Early Treatment Diabetic Retinopathy Study

1.4. MRI data pre-processing

In detail, the following preprocessing steps were applied using each participants' rs-fMRI and structural scan: discarding of first five volumes of the rs-fMRl scan to account for scanner gradient stabilization, realignment to the first volume using b-spline interpolation, slice-timing correction using sinc-interpolation to time-shift and resample functional data slices, tissue segmentation and spatial normalization to standard MNI space including resampling of functional data to 2mm isotropic voxels and structural data to 1mm isotropic voxels, and finally, spatial smoothing by convolving with a Gaussian kernel of 8mm full width at half maximum.

1.5. Group independent comment analysis (GICA)

The GICA feature of the GIFT toolbox was used to reduce all participants' preprocessed rs-fMRI data into 100 group spatial independent components. To this end, each participant's 4D-functional data (255 volumes) were reduced to 150 temporal principal components (PC) that were maximally independent. All participant components were then concatenated, and a second principal component analysis (PCA) was performed on the whole-sample matrix, resulting in 100 PCs. From this PCA matrix, the *Infomax* algorithm [2] for ICA was used to extract 100 group independent components (ICs), which were then checked for stability in *ICASSO* [3] by performing 20 iterations of the ICA algorithm. Lastly, for each participant, a set of 100 component spatial maps and corresponding component time-courses were extracted with the spatiotemporal regression back-reconstruction algorithm [4] in GIFT.

1.6. Component rating and time-course preprocessing

Resulting spatial group independent components were assessed in order to discard components corresponding to physiological, motion-related, or scanner/acquisition-related noise. We considered three main factors during this evaluation as detailed in [5]: (1) location of peak voxels in the corresponding spatial map; (2) peak frequency content in the power spectral density plot and the ratio of high-to-low frequency content; (3) structure of the component timeseries plot. Based on criteria described in [5], the following were characteristics of "noise" components: an IC was classified as "noise" if its spatial map showed clusters that overlapped with regions known to be especially prone to noise or non-neuronal tissue (e.g., white matter, cerebrospinal fluid, vessels), contained many small, distributed clusters, clusters especially near the edges of the brain or clusters which were ring-like in shape. If the power spectra of an IC did not show predominantly low-frequency power or showed a peak or plateau outside of the 0.01-0.1 Hz range, a classification as "noise" was given. Additionally, the IC time-course was inspected for regularity in the oscillation pattern, especially for sudden jumps in the signal which would suggest motion-related noise. Group ICs were manually classified as by two independent raters (AR, NS), and any disagreement was resolved through rating by a third independent rater (CF).

Using the temporal dFNC feature of GIFT, participant's component time-courses corresponding to the remaining "signal" components underwent additional steps of nuisance regression. This included despiking to remove sporadically occurring sharp peaks and detrending to remove mean and linear trends (e.g., frequency signal drifts) as well as motion correction through regression of participant-specific realignment parameters (6 directions) and their derivatives. Finally, the component time-courses were bandpass filtered using a fifth-order Butterworth filter to retain frequencies between 0.01-0.15 Hz. Fully preprocessed time-courses were then read-out from GIFT for later static FC analysis which was performed with custom MATLAB scripts.

An approximate anatomical label for each group IC was determined by visual inspection. Each component was overlaid on a T1-weighted MNI template in the Oxford Centre for Functional Magnetic Resonance Imaging of the Brain Software Library (FSL) tool "FSLeyes". The built-in Harvard-Oxford cortical and subcortical atlases were then used to identify the anatomical label corresponding to peak voxel intensity coordinates of each component. Composite spatial maps grouped by designated resting-state network and anatomical labels for each component are provided in supplemental materials figure 2 and table 2.

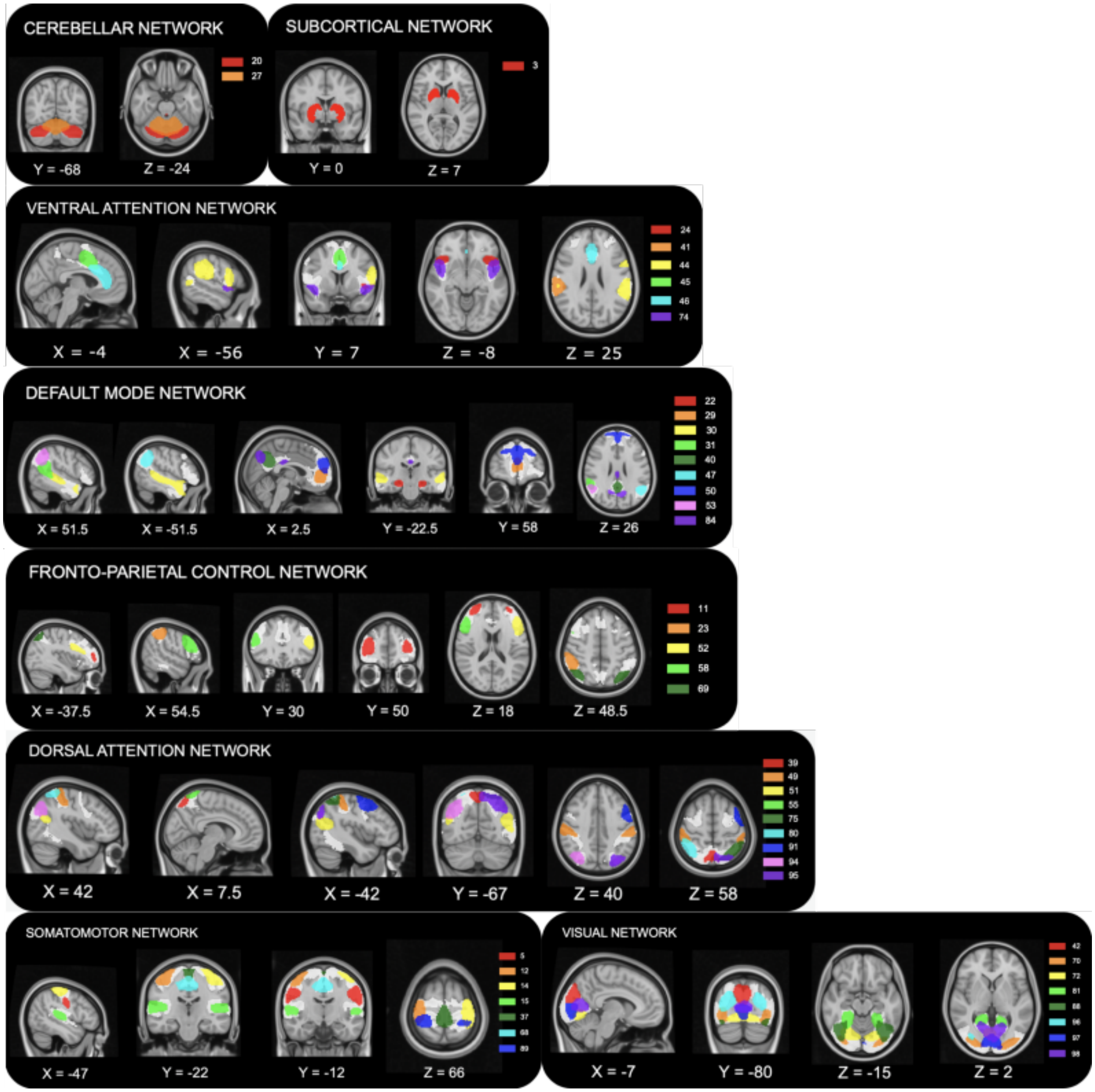


**Supplementary figure 3. Group independent components rated as "signal" and assigned to resting-state networks.** 47 signal components out of 100 components resulting from GICA. Components were assigned to functional networks according to (Yeo et al. 2011), whose binary network masks are shown in white. Each panel corresponds to a functional network and each color corresponds to a component. For visualization purposes, components were thresholded to show only the highest 40% of voxel intensity values. Brain plots are displayed in radiological convention.

**Supplementary table 2 -** Expanded component statistics and brain region labels. Table includes (left to right) component number, brain region label associated with peak voxel coordinates based on the Harvard-Oxford cortical and subcortical structural atlases, maximum voxel intensity, MNI coordinates of peak voxel, and number of voxels in the component.

| | | | | |
|---|---|---|---|---|
| **Ventral attention network** | | | | |
| 24 | Frontal operculum cortex (left) | 8.563 | [-46 18 -4] | 1358 |
| 41 | Supramarginal gyrus, anterior division (right) | 6.628 | [60 -26 26] | 1507 |
| 44 | Supramarginal gyrus, anterior division (left) | 4.518 | [-62 -34 26] | 2663 |
| 45 | Paracingulate gyrus (bil.) | 7.487 | [-2 8 48] | 1707 |
| 46 | Cingulate gyrus, anterior division (bil.) | 5.809 | [-2 32 22] | 2016 |
| 74 | Insular cortex (bil.) | 8.427 | [-46 10 -8] | 1699 |
| **Dorsal attention network** | | | | |
| 39 | Precuneus cortex (bil). | 12.545 | [0 -64 62] | 832 |
| 49 | Supramarginal gyrus, anterior division (bil.) | 5.511 | [54 -22 52] | 2918 |
| 51 | Angular gyrus (bil.) | 5.549 | [-54 -62 18] | 2459 |
| 55 | Postcentral gyrus | 13.211 | [-2 -48 74] | 718 |
| 75 | Superior parietal lobule (left) | 8.565 | [-36 -56 64] | 1280 |
| 80 | Superior parietal lobule (right) | 9.994 | [36 -50 64] | 927 |
| 91 | Middle frontal gyrus (bil.) | 6.659 | [-50 4 54] | 1553 |
| 94 | Lateral occipital cortex, superior division (right) | 6.951 | [38 -70 38] | 1608 |
| 95 | Lateral occipital cortex, superior division (left) | 7.605 | [-28 -72 52] | 1777 |
| **Somatomotor network** | | | | |
| 5 | Precentral gyrus (bil.) | 8.252 | [56 -2 30] | 2208 |
| 12 | Precentral gyrus (right) | 9.161 | [36 -20 70] | 1544 |
| 14 | Precentral gyrus (left) | 8.101 | [-36 -22 72] | 1886 |
| 15 | Planum Temporale (bil.) | 7.171 | [58 -20 14] | 2887 |
| 37 | Precentral gyrus (bil.) | 8.937 | [-2 -32 76] | 1653 |
| 68 | Juxtapositional Lobule Cortex (formerly Supplementary Motor Cortex) (bil.) | 6.500 | [-2 -14 52] | 2241 |
| 89 | Postcentral gyrus (bil.) | 8.683 | [-24 40 76] | 1483 |
| **Visual network** | | | | |
| 42 | Cuneal cortex (bil.) | 7.589 | [0 -80 34] | 2283 |
| 70 | Lateral Occipital Cortex, inferior division (bil.) | 5.933 | [-32 -90 2] | 2668 |
| 72 | Lingual gyrus (bil.) | 6.270 | [8 -70 -6] | 2838 |
| 81 | Lingual gyrus (bil.) | 5.351 | [24 -42-10] | 3222 |
| 88 | Temporal occipital fusiform cortex (bil.) | 6.509 | [-46 -62 -18] | 2329 |
| 96 | Lateral occipital cortex, superior division (bil.) | 4.614 | [30 -76 22] | 3276 |
| 97 | Intracalcarine cortex (bil.) | 8.916 | [0 -86 0] | 1790 |
| 98 | Intracalcarine cortex (bil.) | 6.936 | [-8 -70 12] | 2657 |
| **Default mode network** | | | | |
| 22 | Hippocampus (bil.) | 7.525 | [22 -16 -12] | 1361 |

| | | | | |
|---|---|---|---|---|
| 29 | Frontal pole | 10.404 | [-2 56 2] | 1076 |
| 30 | Superior temporal gyrus, posterior division (bil). | 5.0727 | [60 -28 0] | 4226 |
| 31 | Angular gyrus (right) | 7.040 | [56 -46 16] | 1769 |
| 40 | Precuneus cortex (bil.) | 9.330 | [-2 -54 36] | 1323 |
| 47 | Angular gyrus (left) | 7.119 | [-52 -58 38] | 1838 |
| 50 | Frontal pole | 9.372 | [-2 60 24] | 1266 |
| 53 | Angular gyrus (right) | 7.748 | [50 -54 40] | 1483 |
| 84 | Precuneus cortex (bil.) | 8.750 | [-4 -68 42] | 1659 |
| **Subcortical network** | | | | |
| 3 | Basal ganglia (bil.) | 7.253 | [-26 8 -2] | 2322 |
| **Cerebellar network** | | | | |
| 20 | Cerebellum (bil.) | 4.364 | [-30 -70 -32] | 5660 |
| 27 | Cerebellum (bil.) | 5.488 | [-2 -62 -22] | 3817 |
| **Fronto-parietal control network** | | | | |
| 11 | Frontal pole | 7.942 | [32 62 8] | 1602 |
| 23 | Supramarginal Gyrus, posterior division (right) | 7.665 | [50 -38 56] | 1342 |
| 52 | Middle frontal gyrus (left) | 6.604 | [-52 16 30] | 1945 |
| 58 | Middle frontal gyrus (right) | 6.634 | [50 18 32] | 1871 |
| 69 | Lateral occipital cortex, superior division (bil.) | 8.311 | [-36 -66 56] | 1376 |

##### 1.7. Dynamic functional network connectivity

Additional sliding window correlation parameters set within the GIFT toolbox included: a rectangular window convolved with a Gaussian kernel of s = 3 to create tapered edges; L1 regularization, repeated 10 times to estimate regularization strength (GIFT manual, https://trendscenter.org/trends/software/gift/docs/v4.0b_gica_manual.pdf). L1 regularization in GIFT entails computing inverse covariance matrices with the estimated Lasso penalty parameter using the *graphical lasso* algorithm [6]. Finally, this step is followed by conversion of window-wise covariance matrices to correlation matrices.

We used a window size of 22TRs (49.5s), and a slide length of 1TR (2.25s), following methodological recommendations from [7]. Thus, each participant's time-courses (containing 255 time points) were segmented into sequential windows (95.45% overlap, 233 windows total), and window-wise connectivity matrices were computed, resulting in 233 47x47 dFC matrices for each participant.

##### 1.8. Clustering into temporal connectivity states

Clustering analysis was performed using the "Post-Processing" feature of the Temporal dFNC toolbox in GIFT. Window-wise correlation matrices for all participants were concatenated and used as input for k-means clustering. The k-means clustering algorithm is an unsupervised machine learning algorithm that relies on several user-specified inputs and will cluster a dataset into k-number of groups, regardless of the underlying data structure. Thus, a data-driven approach as implemented in GIFT was used to determine the optimal number of states into which the total group dFNC data would be clustered. So called "cluster estimation" in GIFT is performed by iteratively running the k-means algorithm, specifying a different number of clusters that increases on each iteration. The results of this step are evaluated by plotting a set of cluster validity indices (CVI), which reveal an optimal number of clusters based on the CVIs mathematical solution (e.g., global minimum). After determining the optimal number of clusters for the dataset, by default within the GIFT toolbox the k-means algorithm is run first on the subset of windowed dFNC matrices with the highest standard deviation. The output centroid positions from this step are then used as initiation positions for a second run of k-means clustering using the entire (all participants, all windows) dFNC data as input.

For the most part, default settings for k-means clustering specified in the GIFT toolbox were used in the present study. However, the distance metric was alternatively set to 'city-block' based on evidence from [8] that this distance measure is preferable for high dimensional datasets as opposed to the default Euclidean distance. The number of replicates was set to 10, and the maximum number of iterations to allow for convergence was set to 1000.

#### 1.9. Dynamic metrics

*Modularity*

The community Louvain algorithm as implemented in the Brain Connectivity Toolbox outputs the optimized community-structure statistic and is based on the notion that a network's optimal structure is achieved when it is subdivided into non-overlapping modules, or groups of nodes, such that the number of within-module connections are maximized, and the number of between-module connections are minimized [9].

*Characterizations of connectivity states across all participants*

In addition to group differences in dynamic metrics within each state, we also calculated the effect of state itself on the variance of each dynamic metric. To this end, we used Kruskal-Wallis (KW) omnibus tests where each dynamic metric was treated as a response variable and state as a 5-level explanatory variable. In cases where the null hypothesis of the KW test was rejected, post-hoc Dunn's tests were used to compute pairwise comparisons and adjustment of the false discovery rate was applied. This is an analogous statistical approach as described in the main text for testing the within-state group effect on a dynamic metric.

### 2.0. Results

#### 2.1. State effects on dynamic metrics

In summary, all pairs of states had significantly different average connectivity, as well as fraction time – except for states 4 and 5. State 1 had a significantly higher dwell time than all other states and state 4 had a significantly higher dwell time than state 2, while other comparisons of dwell time were not significant. States 1 and 3 had significantly higher modularity than states 2, 4, and 5, while states 2 and 4 had significantly higher modularity than state 5. Supplementary figure 4 depicts the difference between states for each dynamic metric computed across all participants.

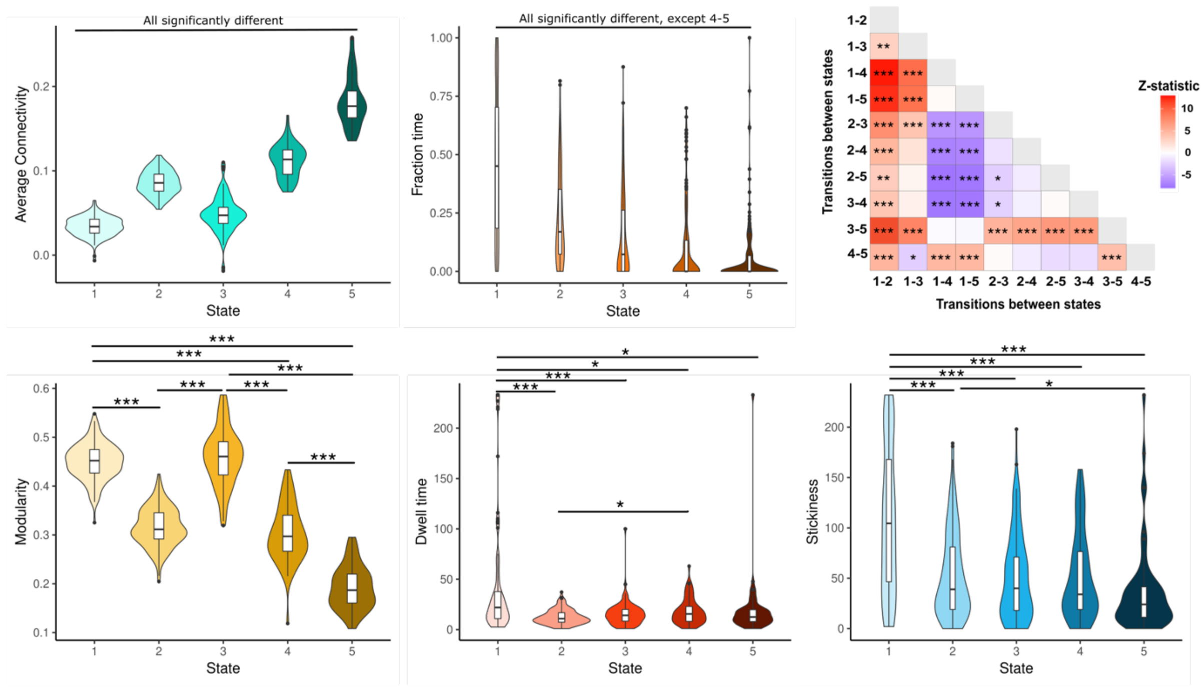


**Supplementary figure 4. State effects on dynamic metrics across the full sample of participants.** Violin plots show state-wise distributions of dynamic metrics across all participant data. Transition matrix (top right) maps the direction and magnitude of Z-statistic and asterisks indicate significance level of differences in number of transitions between each pair of states. * = $p_{fdr} < 0.05$, ** = $p_{fdr} < 0.01$, *** = $p_{fdr} < 0.001$.

#### 2.2. Static network-wise overall connectivity (NWOC)

We performed control analyses using static FC data to compute NWOC to assess whether the inter-NWOC FPN differences and clinical correlations detected with dynamic FC were also a feature of static NWOC. As with dynamic NWOC, no within-network connectivity differences between groups were found. Unlike with dynamic FC, there was no significant group effect on static inter-NWOC between the FPN and the rest of the brain ($c^2 = 5.636$, $p = 0.060$). An insignificant trend in the same direction as seen with dynamic NWOC was found, such that patients with higher disease severity trended towards increased FPN inter-network connectivity compared to less affected patients ($Z = -1.845$, $p_{FDR} = 0.0976$) and HCs ($Z = -2.276$, $p_{FDR} = 0.0691$).

However, to remain consistent with the approach taken in across-state NWOC analyses, we further explored the individual network pairs between FPN and other networks for group effects. Interestingly, as in across-state NWOC calculations, there was a significant group effect on static NWOC between FPN-SMN ($c^2$ = 9.470, p = 0.009) and FPN-SC ($c^2$ = 7.691, p = 0.021). Additionally, a significant group effect on FPN-vATT static NWOC ($c^2$ = 6.084, p = 0.048) was observed – a finding that was not detected using calculations with the dynamic FC matrices.

Multiple pairwise comparisons revealed that patients with EDSS≥2 had a significantly increased FPN-SMN static NWOC compared to patients with EDSS≤1 (Z = -2.744, $p_{FDR}$ = 0.018) and HCs (Z = -2.694, $p_{FDR}$ = 0.011). On the other hand, patients with EDSS≥2 had a significantly decreased FPN-SC static NWOC compared to patients with EDSS≤1 (Z = 2.628, $p_{FDR}$ = 0.026) and HCs (Z = 2.209, $p_{FDR}$ = 0.041). These results were in-line with those of across-state NWOC. We also observed that patients with EDSS≥2 had significantly increased FPN-vATT static NWOC compared to HCs only (Z = -2.457, $p_{FDR}$ = 0.042).
